# Dissociating the Neural Correlates of Planning and Executing Tasks with Nested Task Sets

**DOI:** 10.1101/2021.08.26.457791

**Authors:** Savannah L. Cookson, Eric H. Schumacher

## Abstract

Task processing (e.g., the preparation and execution of responses) and task representation (e.g., the activation and maintenance of stimulus-response and context information) are two facets of cognitive control supported by lateral frontal cortex (LFC). However, the mechanistic overlap (or distinction) between these two facets is unknown. We explored this by combining a complex task mapping with a pre-cueing procedure. Participants made match/non-match judgments on pairs of stimuli during fMRI recording. Pre-cues on each trial gave variable amounts of information to the participant in anticipation of the stimulus. Our results demonstrated that regions throughout LFC were more active at the stimulus (when responses could be executed) than at the cue (when they could only be prepared), indicating that they supported execution of the task agnostic to the specific task representation. A subset of regions in left caudal LFC showed increased activity with more cue information at the cue and the reverse at the stimulus, suggesting their involvement in reducing uncertainty within the task representation. These results suggest that one component of task processing is preparing and executing the task according to the relevant representation, confined to left caudal LFC, while non-representational functions that occur primarily during execution are supported by different regions throughout the rest of LFC. We further conducted an exploratory investigation of connectivity between the two groups of regions in this study and their potential relationship to the fronto-parietal and cingulo-opercular networks. Regions with both patterns of activity appear to be part of the fronto-parietal network.

---

Cognitive control refers to the set of psychological processes and representations that allow us to formulate goals based on our current internal mental state and to use those goals to guide our perception of and interactions with the external world. Two intertwining aspects of cognitive control are our capacity to 1) rapidly and flexibly construct a mental framework of a given task, i.e., task representation; and 2) to use information in the environment to prepare for and execute that task, i.e., task processing. While both of these facets of cognitive control are known to be supported in part by regions in the lateral frontal cortex (LFC; Badre & D’Esposito, 2009; Schumacher et al., 2007; Waskom & Wagner, 2017), their literatures are largely separate. Research on task processing has shown that we can use information about what aspects of a task are currently relevant to select and prepare a subset of that stimuli and/or responses a priori, and that this preparation is supported by LFC regions that are largely distinct from those involved in executing the response (Cookson et al., 2016; Dixon et al., 2014). In parallel, research on the representation of tasks has revealed that we represent tasks with a multi-level nested structure, abstracting out high level contextual and episodic task components that can be selected for to then guide selection between simple stimulus-response associations (Hazeltine & Schumacher, 2016; Schumacher & Hazeltine, 2016). Much like the distinct LFC contributions to task preparation and execution, different LFC regions organized along a rostro-caudal axis are thought to represent progressively more abstract levels of the task within this nested structure (Badre, 2008; Koechlin & Summerfield, 2007; Nee & D’Esposito, 2016). Here, we present a novel task that aims to investigate the LFC correlates of preparation and execution, two components of task processing, at distinct levels of abstraction, a key facet of task representation.

There is a long and broad history of research on the psychological and neurological mechanisms for task preparation and execution. A common procedure for investigating these mechanisms is the pre-cueing procedure, which presents information about a task sequentially at two different times that can be statistically separated by way of a variable delay period. This separation in time allows researchers to distinguish the processes that select and prepare a response from those involved in executing that response. Early work with the pre-cueing paradigm, focused on motor selection and preparation, demonstrated that participants could prepare subsets of motor responses prior to having full knowledge of the required response, which resulted in a response time (RT) benefit for informative relative to uninformative cues (Adam et al., 2003; Miller, 1982, 1985; Reeve & Proctor, 1984). The results of this work suggests that participants were not waiting until all information was known to begin specifying components of the upcoming response, as would be implied by the fully serial processing models common at the time; rather, they were preparing subcomponents of the response – e.g., response hand, movement dimensions, etc. – as the information given allowed them to do so.

Hopfinger, Buonocore, and Mangun (2000) used event-related functional magnetic resonance imaging (fMRI) to investigate whether the regions involved in these preparation processes had distinct neural bases from those that supported execution of that response. On each trial of the task, the authors presented a directional followed by a pair of horizontally aligned checkerboard stimuli. Participants then made a discrimination decision on the checkerboard presented on the cued side. The results showed marked differences in the activation patterns between the preparation and execution phases of the experiment. This distinction was especially apparent in LFC, where preparation showed widespread left-lateralized frontal activation in superior and middle frontal gyri, while execution showed a more constrained left-lateralized activation of inferior frontal gyrus coupled with bilateral activation of the motor cortices. Thus, preparation of a response appears to be mediated by largely distinct brain regions from those supporting execution once the full response is known.

Beyond simple distinctions between preparation and execution, research using the pre-cueing procedure further aimed to understand the source of the cue benefit itself. Rosenbaum (1980) first demonstrated that the magnitude of the behavioral benefit afforded by a cue was dependent on the number of distinct pieces of information given about the upcoming response, independent of the type of information given. Participants were instructed to make movements specified by three independent dimensions: hand (left/right), direction (toward/away from the body), and extent (short/long). They were then presented a cue on each trial that could indicate one, two, all three, or none of the values of the upcoming movement along these three dimensions. The RTs showed a main effect of the number of dimensions remaining to be specified at the time of stimulus presentation. That is, participants responded faster the more dimensions that were cued prior to the time of response. Later research using positron emission tomography demonstrated that this cue benefit likewise was associated with the LFC (Deiber et al., 1996). This suggests that participants made use of the cues to prepare sets of potential movements given whatever partial information they received about the upcoming response.

Evidence shows that, under certain situations, participants can benefit from cue information that indicates one subgroup of stimuli and responses over another, even when that cue information does not confer explicit information about the upcoming response. (Adam et al., 2003). This suggests that preparation is not confined to low-level responses, but that participants can select between whole subsets of a task, or “task sets” (Sakai, 2008). Recent work from our group has demonstrated a pre-cueing benefit specifically for when a task set is cued regardless of the response set (Cookson et al., 2019). In this design, participants learned stimulus-response mappings that either segregated different stimulus categories to different hands (”Separated”) or interleaved them across the fingers (”Interleaved”). Cues were then given that could indicate the upcoming stimulus category (Face or Scene images) or response hand (Left or Right). The group that learned the Separated mapping showed a larger benefit of cue information than the group that learned an Interleaved mapping, suggesting a benefit of preparing a coherent task set above and beyond the benefit of information that simply restricted the total number of potential stimulus-response options. Notably, this was even true for response hand cues, suggesting an additional benefit of selecting an abstract task set above and beyond the benefits of cues giving direct motor response information. In other words, there appear to be distinct selection mechanisms operating at different abstraction levels of a task’s representational structure.

Though interest in the cue benefit originated from research on task processing, it has since become a cornerstone of the task representation literature. The idea that participants can select between subsets of stimulus-response associations, and even subsets of subsets, is formalized in the task file hypothesis, which proposes that task processing is guided and bounded by the structure of our internal representation of a task’s structure. An extension of the event file hypothesis (Hommel, 2004), The task file hypothesis posits that tasks are represented as associations of stimuli, responses, task contexts, and other goal-related factors (Schumacher & Hazeltine, 2016). These task files are highly flexible, allowing stimulus-response associations to be rapidly grouped together into meaningful subsets bound to distinct contextual triggers according to current task context and selected between on demand. Task sets can be further extrapolated to more abstracted structures that add nested layers of additional context associations, allowing for the selection between entire task sets as a whole.

Work on understanding how nested task set representations support and bound task processing has remained largely behavioral thus far. However, investigations of the role of LFC regions in cognitive control outside of the task set literature suggest that the division of task representations in the brain follows a similarly nested structure (Badre, 2008). The areas of activation seen in these studies are strongly left-lateralized and are connected to one another along a rostro-caudal axis (Badre & D’Esposito, 2007; Koechlin & Summerfield, 2007). These regions appear to represent increasingly abstract aspects of a task progressing from caudal to rostral regions. Caudal areas represent the task-relevant, concrete associations between stimuli and responses; mid-frontal areas represent context-dependent rules; and rostral areas maintain schema that organize and contextualize the set of potential task rules and action sequences that apply to the task (Badre & Nee, 2018).

Studies relating LFC regions to different levels of task abstraction often use tasks that manipulate the number of levels in a complex response mapping structure. Badre and D’Esposito (2007; see also Koechlin & Summerfield, 2007 for an alternative hypothesis with similar design and results) reported one such task that distinguished between four levels of abstraction. At the lowest level, participants selected responses to presented stimuli (colored squares) based on a learned mapping. The next level used cue-dependent responses, in which participants might have to respond to one of a number of potential target features based on the cue presented (e.g., respond to the texture of a target when the cue color was red or to its orientation when the cue color was blue). At the third level, participants were shown pairs of stimuli and instructed to make match/non-match decisions on the feature dimension indicated by the cue, rather than making a response based on the value of the feature itself. For example, participants might have to decide if the textures of two stimuli were the same when the cue color was red, and if they were at the same orientation angle when the cue color is blue. Finally, the fourth level implemented the same third level task, but with block-level switching of the mappings between cues and dimensions. At each level of abstraction, then, the design added another layer of contingencies that had to be resolved in order to correctly execute the task. Functional magnetic resonance imaging (fMRI) recordings taken during the execution of this task demonstrated progressively more rostral activation in LFC for tasks at higher levels of abstraction.

The multiple layers of contingencies used in the task above are similar to the nested task set structures described in the task file hypothesis (Hazeltine & Schumacher, 2016; Schumacher & Hazeltine, 2016). However, because imaging studies investigating the frontal organization of cognitive control have mainly relied on trial-level or mixed-block designs, little is known about how preparation and execution may be instantiated in the brain across distinct levels of representation. Conversely, imaging studies investigating task preparation and execution are limited and have mainly relied on simple stimulus and motor cues rather than the abstract cues that are used in task set pre-cueing. In another study from our group (Cookson et al., 2016), we explored the neural correlates of preparation and execution for cued and uncued task sets using the same “Separated” mapping task described above (viz., Cookson et al., 2019) and presenting cues that either gave information about the upcoming stimulus category or response hand (”informative”) or neither (”noninformative”). In this way, when participants were given an informative cue, it indicated one set or the other for the upcoming stimulus, again without necessarily imparting explicit motor information. The noninformative cue, on the other hand, would indicate that either set could be relevant. fMRI data recorded during the presentation of the cue showed greater LFC activity to informative cues of both types relative to noninformative cues, suggesting that participants prepared the individual task sets at the cue when possible. Alternatively, this may have also been an indication that participants were processing the information contained in the cue to determine and prepare the implied motor effector. While our previous behavioral work (i.e., Cookson et al., 2019) suggests that task set selection can benefit performance above and beyond motor preparation, this design could not completely disentangle the two. Thus, it is still not clear how cue-related preparation and subsequent execution may be instantiated at different levels of task abstraction.

Thus, both preparation and execution, components of task processing, and task abstraction, a facet of task representation, have similar psychological constructs and both show distinct contributions of different regions of LFC over the course of the task. However, it is still unknown whether and to what extent the mechanisms involved in these aspects of processing and representation are shared or distinct. If different sets of brain regions support distinct psychological mechanisms of task processing and representation, two distinct subsets of task-related regions should emerge. Regions mediating task preparation and execution should show general differences in activation between the cue and stimulus time point; specifically, regions mediating preparation should activate to the cue whereas regions mediating execution should activate to the stimulus. Regions mediating task representation may show activity that is independent of time (i.e., independent of cue versus stimulus phase during the trial) but whose activity varies with the amount of task information available. This may emerge as a distinction between cue-sensitivity in the LFC regions previously identified by policy abstraction studies (see review by Badre, 2008) and time sensitivity (i.e., differential cue and stimulus phase activity) in other task-related regions, such as motor cortex and SMA. On the other hand, task representations may be the scaffolding on which time-dependent processing takes place. In this hypothesis, some or all LFC regions may show sensitivity to the amount of information provided by the cue information, but this sensitivity may also depend on the time or phase of the trial. This variation in time may depend on the amount of information being processed simultaneously as the system works to reduce uncertainty about the response as information is made available, or may vary with the amount of information left to specify, indicating that they are representing a larger potential task set. Variations as a function of cue information may further depend on a given region’s position along the rostro-caudal axis, with more rostral regions being more sensitive to abstract task information and more caudal regions being related to low-level stimulus-response information.

Here, we present a novel task that blends a pre-cueing paradigm with a pseudo-hierarchical response mapping structure to investigate the neural correlates of preparation and execution for multiple levels of task representation, illustrated in Figure 1. During the experiment, participants were presented with pairs of stimuli and instructed to make a match/non-match judgment on one of four possible dimensions. The judgment domains could be categorized into two groups: two of the judgment dimensions related to spatial features of the stimuli, and two to object identity (i.e., non-spatial) features. Each judgment dimension was associated with a two-choice response mapping. The responses for one judgment dimension from each domain (i.e., spatial and non-spatial) were mapped to each hand, where each possible response was mapped to a different response finger. At the start of each trial, a cue was presented that gave participants information about either the upcoming domain of the relevant judgment dimension, the hand with which participants would make their response, both, or neither. Because judgment domain and response hand were crossed in this design, information about either could not be used to determine the other, though each would allow participants to prepare a subset of half the task structure; that is, while both cues narrowed the response set the same amount, only cues for response hand allowed for specification of any motor dimensions. Information about both would implicitly specify the judgment dimension for the upcoming trial, allowing participants to prepare for a 2-alternative forced choice task. This allowed us to explore preparation versus execution in three levels of cue information.

**Figure 1.**
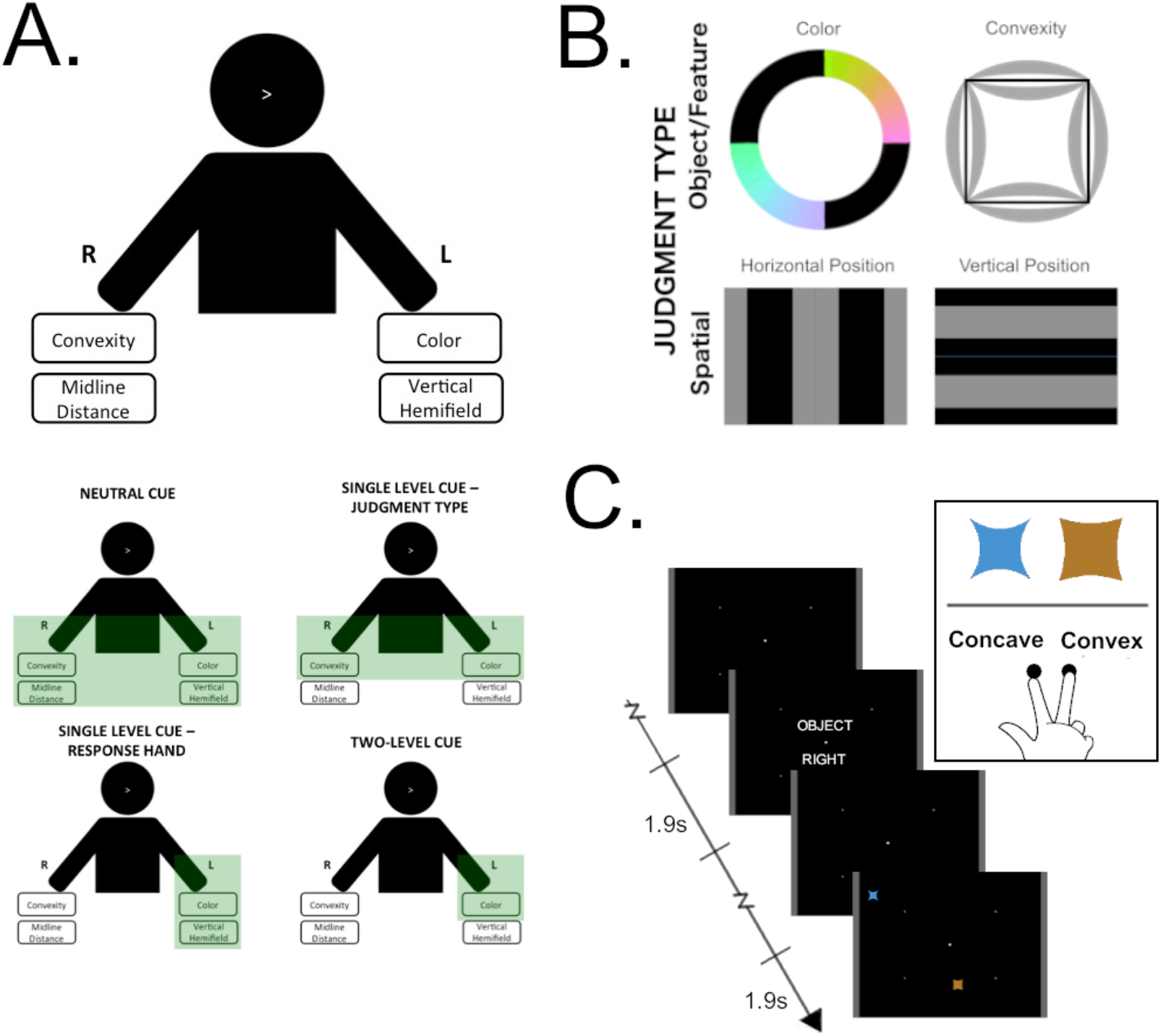
Illustration of experimental design elements. A) Participants learned a task mapping that mapped one each of two spatial (vertical hemifield and midline distance) and two object-based judgments (convexity and color) to each hand. Cues presented at the start of each trial could provide information about either the upcoming judgment dimension, response hand, both, or neither, which reduced the set to the subset highlighted in green for each cue. Cues with no information left the full task set remaining to be specified; cues for the judgment domain reduced the set by half but did not reduce the decision to a single hand; cues for response hand limited the set to one hand and reduced it by half; and cues for both pieces of information reduced the set to a forced choice task. B) Visual ranges of the four features that defined the stimulus pairs on each trial. Features were selected from two ranges, shown in grey (for color, shown as non-blacked out subsets of the DKL color wheel), that corresponded to different judgment decisions for that feature. C) Example trial. Participants saw a cue following a varied inter-trial interval for 1.9s. They then waited for a variable cue-stimulus interval before the stimulus pair for that trial was presented. Participants determined which of the four features of the stimulus pair shared the same judgment decision for both stimuli – in this case, both stimuli are convex – and then make a response for that judgment decision based on their learned mapping.

Using this design, we defined a set of LFC regions that were involved in the task as a whole (see Methods for details), then assessed how each region’s activity varied between the cue and stimulus time points of each trial as a function of the amount of information contained in the cue. Regions that were differentially involved in preparation and execution should demonstrate activity that varied between the cue and stimulus presentations. Specifically, regions mediating preparation will activate to the cue, while regions mediating execution will activate to the stimulus. Regions that manage representations of the task should show sensitivity to the amount of cue information as they prepare different subcomponents of the task on each trial. This design allowed us to test several key questions surrounding the relationship between preparation and execution and task abstraction.

First, we asked whether distinct regions were involved in preparation and execution versus task abstraction, or if some or all regions would have overlapping involvement in both. If they are supported by distinct mechanisms, regions should demonstrate sensitivity to the time point in the trial or the level of cue information, but not both. Conversely, if they share underlying mechanisms, some or all of the regions in our task should demonstrate an interaction effect between time point and cue information. This analysis would also allow us to explore whether cue-sensitive regions showed distinct activity patterns for cues that imparted motor information versus those that did not.

Second, we asked how different amounts of cue information might drive activity levels in cue-sensitive regions. If cue sensitive activity is the result of the preparation of a relevant task set, or alternatively a marker of the number of alternative potential responses being prepared, regions showing this cue sensitivity should have greater activity as a function of the size of the set made relevant at the cue. For example, for a noninformative cue, no stimuli or responses can be excluded and thus the full task mapping remains relevant. Thus, we would expect the greatest activity for these cues. On the other hand, a cue for both judgment domain and response hand can reduce the task to a simple two-alternative forced-choice decision. In this case, we would expect the least amount of activity, with cues for one or the other in between. As an alternative hypothesis, it is possible that activity in LFC regions represents the resolution of uncertainty as information becomes available. In this case, we would expect the reverse pattern; cues giving both the domain and response hand would allow for the selection of two levels of nested task sets (i.e., reducing two points of uncertainty in the task mapping structure), and would therefore show the most activation at the cue, whereas noninformative cues provide no resolution of uncertainty and would show minimal activity. Moreover, this alternative hypothesis further predicts a reversal of this pattern at the stimulus, at which point the remaining information of the task can be processed. Following this investigation, we further asked how these patterns were distributed along the rostro-caudal axis in our left-lateralized frontal regions.

## Methods

### Participants

Twenty right-handed participants (10 female) ages 18-34 were recruited from the student population at the Georgia Institute of Technology. This sample size was selected to be consistent with the median sample size of fMRI studies at the time of collection (Poldrack et al., 2017). One participant was removed from analysis due to a technical issue with data collection and one participant was removed due to sub-criterion performance on session 2. Participants had normal or corrected-to-normal vision, and were not otherwise contra-indicated for an MRI protocol. Participants gave written informed consent under the Georgia Institute of Technology Institutional Review Board and were compensated with course credit.

### Apparatus

Stimuli were presented using PsychoPy software (Peirce, 2007) via a personal computer (PC) connected to an Avotec Projector (mock and MRI scanners). Task code is available at http://www.github.com/savannahcookson/Hierarchical-Precue-Task. Responses were collected using two scanner-compatible PST button boxes. Participants wore covered protective earphones in both sessions, with earplugs in the MRI for additional protection from noise. In the MRI, head movement was restricted with soft padding. In the mock scanner, scanner noises were simulated using a standard CD player with sounds recorded during typical MRI sessions. All presentation, scanning, and response collection apparatus were available through the Georgia State University/Georgia Institute of Technology Center for Advanced Brain Imaging.

The screen was presented at a resolution of 1024×768px with a black background. Text, including the instructions, cues, feedback, and fixation screens, was presented in white font. To provide a fixation point and additional spatial reference points for the task decisions, a square marker (80px) was shown in the center of the screen. This marker was surrounded by four smaller (40px) square points arranged in a rectangle centered around the fixation point such that each of the four points marked a corner of the rectangle. Each of these four points was exactly halfway between the center and outer edges of the screen in both directions. Grey bars were shown on the left and right edges of the screen to delineate the outer bounds of the horizontal aspect of the screen. Screen center is defined as point (0,0) for pixel ranges.

### Target stimuli and cues

Stimuli consisted of warped, colored squares presented in locations that varied both vertically and horizontally (Figure 1.B). Descriptions of each of these four features, and the judgments participants made for each, are described below; precise value ranges for each of the four features of the stimuli are presented in Table 1.

**Table 1.**
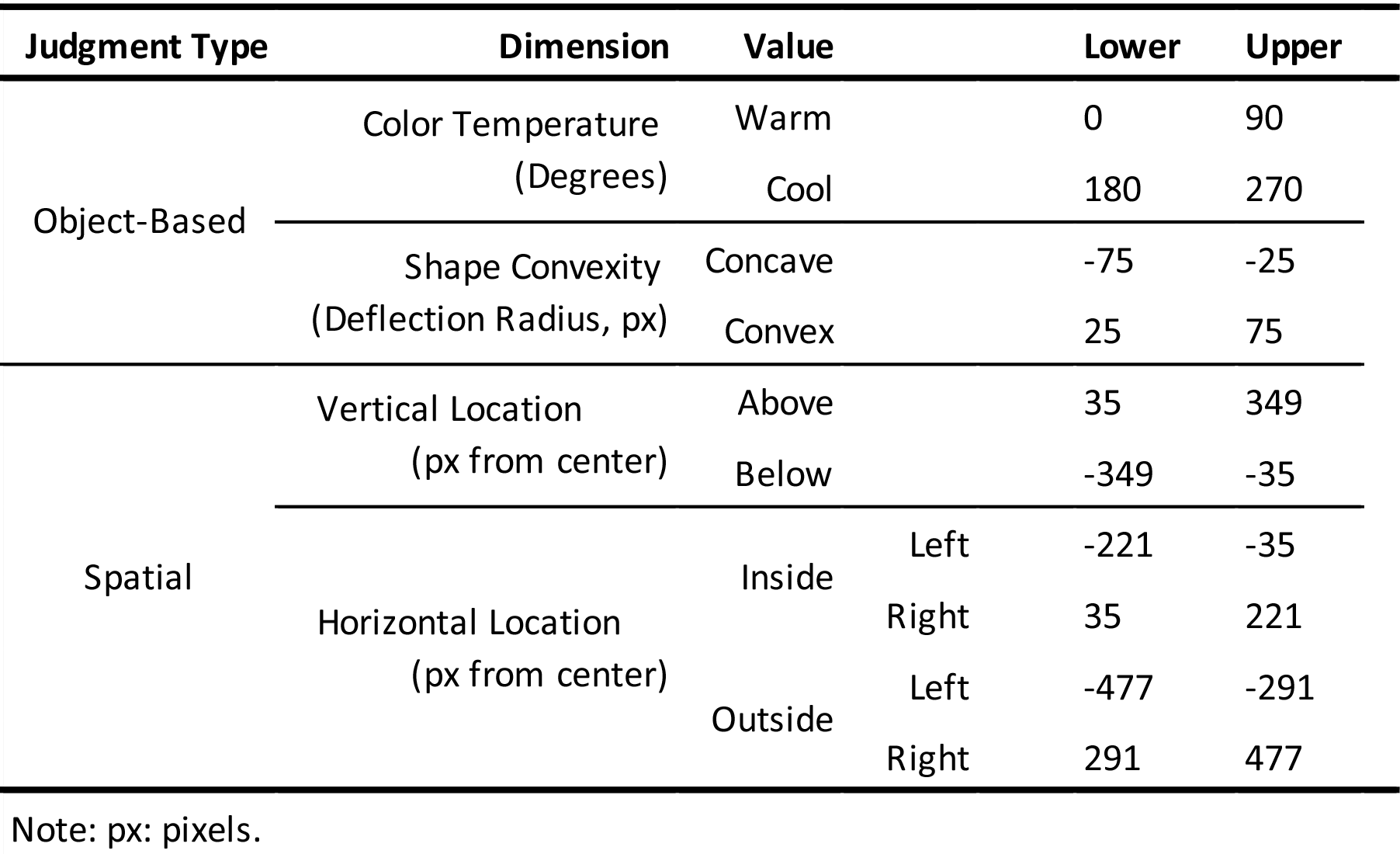
Value ranges for stimulus features by judgment type.

#### Color temperature

The color of each stimulus was defined along Derrington-Krauskopf-Lennie (DKL; Derrington et al., 1984) color space. In this space, the range of observable colors are plotted as a color wheel oriented orthogonally to the luminance of the observed color; here, luminance was held constant at 0° to approximate isoluminance across our color ranges. Degree ranges along this plane were set for “warm” and “cool” color judgments such that “warm” colors include reds, oranges, yellows, and a limited range of yellow-greens; “cool” colors, likewise, include a subset of dark greens, blues, and purples.

Shape convexity. The shape of each stimulus was derived from a square 200 pixels (px) on each side. The sides of the template square were warped so that the center of each line was deflected toward or away from the center to form a quadratic parametric curve with the corners of the square maintained in their original positions. The deflection range in each direction varied with a resolution of 1px. Such warping allowed participants to judge the shape as a “concave” (deflected inward) or “convex” (deflected outward) shape. These stimulus images were resampled to a size of 70px square for experiment presentation.

#### Vertical location

The vertical location dimension established the y-coordinate of the presentation location for each stimulus, independent of the horizontal location (x-coordinate). Vertical location was defined as a function of the height of the presentation window (1024px). Stimuli could be presented at a range of distances from the horizontal midline of the screen with resolution of 1px, where stimulus position was determined by the location of the center point; the vertical location of the stimulus could then be judged as “above” or “below” the midline. Horizontal location. Horizontal location was defined as a function of the width of the presentation window (1280px). Stimuli could be presented at a range of distances from the vertical midline with a resolution of 1px, again determined by the location of the center point, with discrete windows in the outer and inner halves of the screen; stimuli could be judged as appearing toward the “inside” or “outside” of the screen.

#### Cues

Cues consisted of two words or letter blocks presented in capital letters, with one presented above and one below the central fixation (Figure 1.C). Cues for judgment domain (”SPATIAL” or “OBJECT”) appeared above the center fixation, and cues for response hand (”LEFT” or “RIGHT”) below. When information was not to be given for a particular cue type on a given trial, “XXXXX” served as the placeholder in the relevant cue position. For example, for a trial in which no information was given, “XXXXX” would be presented both above and below the center fixation. For a trial in which only judgment domain was indicated, “SPATIAL” or “OBJECT” would be presented above fixation as appropriate, with “XXXXX” shown below; conversely, for trials that only indicated response hand, “XXXXX” was shown above fixation, with “LEFT” or “RIGHT” shown below as needed. In trials in which both pieces of information were given, the relevant judgement type was shown above and the relevant response hand below, with no “XXXXX” presented in either position.

### Procedure

The experiment proceeded in two sessions; the first session was completed in the mock scanner, and the second in the MRI scanner. When participants arrived for the first session of the experiment, they completed the informed consent process and were assigned to a mapping structure. In each mapping, one dimension from each judgment domain (object, spatial) was mapped to each hand; i.e., one object-based and one spatial judgment each were mapped to each hand (Figure 1.A). Index and middle finger mappings for judgments were assigned for each of the two decisional options of the mapped dimension. These mappings were partially counterbalanced between participants for a total of 8 mapping sets. Participants were trained on their assigned mapping at the start of session 1, as described below, and complete session 2 using the same mapping.

Once the consent process was complete in session 1, the participant was oriented to and assisted into the mock scanner. They then began the training phase of the task, which took approximately 30 minutes. Next, the participant completed between 6 and 12 experimental runs. The number of runs completed by each participant was determined by the how quickly the participant reaches a criterion performance; specifically, an average accuracy of 90% over the three most recent blocks. At the start of the first of these runs, the participant was informed that the experimenter would start playing the MRI sounds, and that they should remain as still as possible. The MRI sounds were played for the duration of each experimental run, and were stopped between runs while the experimenter checked in with the participant.

The second session took place 1-7 days after the first. Upon arrival, participants were assisted into the MRI scanner. Once the participant was ready to begin, they completed a shortened version of the training procedure described below. Finally, they completed 16 experimental runs. The experimenter checked in with the participant briefly in between runs. Upon completion of the task, participants were extracted from the scanner and brought back to the lab, where they completed a debriefing questionnaire.

Experimental runs in both sessions consisted of 32 trials. Runs presented all possible combinations of cue types, judgment domains, response hands, and decision values. In both sessions, participants were given overall feedback for their accuracy and RT across the run at the end of that run. In addition, during session 1, feedback was given at the end of each trial. If participants gave the correct response, they were shown the fixation screen; if they gave an incorrect response, they were told what the correct response should have been.

#### Training

To learn the task, participants completed a series of progressive training blocks in session 1. First, participants familiarized themselves to the four possible responses in the task (index or middle finger press; the left or right hand) by responding to explicit instructions, e.g., “Press the RIGHT MIDDLE finger.” This practice block contained 8 trials, counterbalanced across the four possible response options.

Next, participants completed a set of mapping training blocks that progressively introduced the mapping for each judgment dimension. These blocks instructed participants on the responses required for each of the four judgment dimensions based on their assigned mapping. At the start of each block, participants were presented with instructions that indicated: 1) what judgment was to be made for that particular dimension; 2) the hand that would be used to respond to that dimension; and 3) what responses corresponded to each of the two possible judgment decisions. Following the instructions for a given dimension, participants completed 16 practice trials for that dimension, in which they performed the instructed judgments on a single stimulus. In all participants, the mapping blocks introduced the dimensions in the following order: color temperature, shape convexity, vertical location, and lastly horizontal location. For the color temperature training block squares presented in the center of the screen bearing a particular color. As participants progressed through the mapping training phase, the stimuli gained values along each of the new dimensions while continuing to present with random values along the already-learned dimensions. This allowed participants to incrementally familiarize themselves with the nature of the stimuli they would see on-screen. When the vertical location dimension was introduced, participants were further instructed to maintain fixation on the center of the screen at all times during the task. The instruction to maintain fixation on the center of the screen was emphasized at the beginning of each subsequent block and run.

Once the mapping training was complete, participants were introduced to the experimental task, without cues. They were instructed that they would see stimuli appear in pairs, and that these pairs of stimuli would share a judgment decision value along exactly one of the four possible dimensions. They were then instructed to identify the shared dimension and respond with the appropriate response for the judgment decision value shared by the pair, according to their previously learned mapping. They then completed a practice block consisting of 32 trials in which they respond to pairs of stimuli as they were presented onscreen. Trials were counterbalanced across judgment domains, response hands, and judgment decision values.

Finally, participants were introduced to the pre-cueing aspect of the task. They were instructed that each trial would be preceded by a cue that may give some information about the upcoming trial. The instructions explained the information that could be contained in the two parts of the cue and describe how these two parts could combine into the different cue levels (no information, hand-only, judgment-only, or both). Participants were instructed to use the cues to the best of their ability to help their performance during the task. Following these instructions, participants completed a full practice run of the experimental procedure, as described below.

#### Trial structure

On each trial, participants made judgments of pairs of stimuli based on the feature that the two stimuli shared. First, the cue was presented for 1.9s. This was followed by a fixation screen for a cue-stimulus interval (CSI) that was jittered as described below. Following this interval, the two stimuli were presented simultaneously on the screen, one in the left half and one in the right, for 1.9s. The stimuli had different judgment values for each of the four dimensions described above except for one; participants were instructed to identify this shared dimension and to make the appropriate index or middle finger response on the hand associated with that judgment decision value. Participants were able to make their response for the interval that the stimuli were on-screen; this was followed by a fixation screen with a inter-trial interval (ITI) that was jittered in the same manner as the CSI. Participants were instructed to respond as quickly and accurately as possible on each trial, and to use the cues presented at the beginning of each trial to prepare the possible upcoming responses to the best of their ability. An illustration of the trial structure is found in Figure 1.C.

The CSI and ITI durations were jittered using the Analysis of Functional NeuroImages (AFNI; Cox, 1996) optimization algorithm. This algorithm produces a random experimental timing given the desired trial types (i.e., the factor combinations for the cue and stimulus time points), the desired average duration of the CSI and ITI events (1.9s each), and a set of bounding parameters (minimum duration: .25s; step: .25s.). An estimate of the unexplained variance for this design can then be calculated. Here, for the experimental runs in each session, this process was iterated 1000 times, comparing the unexplained variance for each design and keeping the design that minimized this value. The same process was completed over 100 iterations to produce the timings for cue practice blocks in the training.

#### fMRI parameters

Images were collected on a Siemens 3T MRI scanner using a standard 32-channel radio-frequency head coil. At the beginning of the session, a 3D MPRAGE structural scan (1mm isotropic voxels) was collected, followed by a three-plane localizer. The blood-oxygen level-dependent (BOLD) signal was recorded during each experimental run using an interleaved echoplanar T2* sequence (TR=2000ms, TE=30ms, 3mm isotropic voxels). Each functional volume contained 37 axial slices, with 130 volumes/run.

### Analyses

#### Behavioral analyses

RT values included correct trials only. Mean RTs and arcsine-transformed accuracies on the experimental blocks for session 2 were calculated for each subject as a function of three within-subjects factors (cue type, 4 cells; judgment domain, 2 cells; response hand, 2 cells) as well as a between-subjects factor for the congruency of each participant’s horizontal judgment response mapping (2 cells; viz, whether the “inner” judgment was mapped to the index or middle finger). This latter factor was included to assess a potential confound of congruency effects due to participants’ natural tendency to map fingers closer to the body to stimuli presented closer to the midline of a display. A four-way repeated measures ANOVA was performed on the RT and accuracy data for the three factors. Post-hoc comparisons were conducted using Tukey’s HSD procedure.

#### fMRI processing and whole-brain analysis

Data preprocessing and analysis were conducted using AFNI (Cox, 1996). Preprocessing procedures included despiking, slice timing, volume registration, 6-parameter rigid-body motion correction, structural-functional alignment and registration to standard Montreal Neurological Institute (MNI) space. The data were scaled to an intensity range of 0 to 100 to reflect percent signal change.

Data were analyzed using typical general linear modeling techniques. Nuisance regressors were included for constant, linear, and quadratic trends, as well as the 6 motion parameters extracted from preprocessing. Events were defined for the cue and stimulus time points (1.9s each) of each trial, labeled for the cue type, judgment domain, and response hand. Beta values were scaled from 0-100 to reflect percent signal change (PSC). Contrasts were then defined for all effects of interest, including task (all cue and stimulus time points) versus baseline.

#### Region of interest definition and activation analysis

To define our regions of interest (ROIs) for analysis, we started by generating a set of 6mm-radius spherical regions across the brain using the coordinates published by Power and colleagues (2011). We then masked this set of regions to the FDR-corrected (q = .05) whole-brain contrast for task versus baseline, restricting final inclusion to those regions that retained at least 10 voxels after thresholding. Average PSC for each within-subjects condition (i.e., cue type, 4 cells; judgment domain, 4 cells; response hand, 2 cells) at each time point (i.e., cue or stimulus event) were extracted for each ROI and each participant. The average PSC values for each condition were then subjected to a four-way repeated measures ANOVA within each region across subjects. Where necessary, violations of the sphericity assumption were corrected for using the Hyunh-Feldt method.

## Results

### Behavioral results

Accuracies and RTs are presented in full in Table 2 as a function of condition.

**Table 2.** Behavioral results across subjects summarized by condition

| Cue Type | Judgment | Hand | Congruent Mapping |  | Incongruent Mapping |  |
| --- | --- | --- | --- | --- | --- | --- |
|  |  |  | Accuracy | RT (ms) | Accuracy | RT (ms) |
| None | Object-Based | Left | 0.95 | 973 | 0.92 | 941 |
|  |  | Right | 0.95 | 930 | 0.94 | 901 |
|  | Spatial | Left | 0.90 | 939 | 0.92 | 974 |
|  |  | Right | 0.93 | 971 | 0.91 | 993 |
| Hand | Object-Based | Left | 0.96 | 882 | 0.93 | 915 |
|  |  | Right | 0.98 | 832 | 0.97 | 856 |
|  | Spatial | Left | 0.98 | 874 | 0.94 | 893 |
|  |  | Right | 0.96 | 874 | 0.94 | 924 |
| Judgment | Object-Based | Left | 0.95 | 906 | 0.93 | 899 |
|  |  | Right | 0.96 | 845 | 0.96 | 856 |
|  | Spatial | Left | 0.93 | 907 | 0.92 | 912 |
|  |  | Right | 0.94 | 890 | 0.95 | 959 |
| Both | Object-Based | Left | 0.98 | 740 | 0.98 | 779 |
|  |  | Right | 0.97 | 712 | 0.97 | 785 |
|  | Spatial | Left | 0.98 | 698 | 0.98 | 733 |
|  |  | Right | 0.97 | 701 | 0.99 | 783 |

Note: Within-subject factors are presented by row; between subject factors are presented in separate columns. RT: reaction time. ms: milliseconds.

#### Accuracy

Accuracies approached ceiling (Overall average = 95.1%). The ANOVA revealed a main effect of cue type, *F*(3, 48) = 15.37, *p* < .001, *η_p_^2^* = .49; in general, participants were more accurate when cues provided more information about the upcoming trial. There was also a significant interaction between cue type and judgment domain, *F*(3, 48) = 2.83, *p* = .048, *η_p_^2^* = .15; that is, the cue-related performance differences above varied based on whether participants were making a spatial or object-based judgment. No other effects for accuracy were significant.

#### Reaction time

The ANOVA revealed a main effect of cue type, *F*(3, 48) = 307.04, *p* < .001, *η_p_^2^* = .95; in general, participants responded faster the more information the cue provided, illustrated in Figure 2. Post-hoc t-tests, summarized in Table 3, confirmed that participants responded fastest in trials giving both judgment domain and response hand information and slowest in trials giving no information. They also showed no significant difference between RTs between single-level cues for response hand and judgment domain, indicating that participants prepared both subsets of the task equally whether a motor response could be prepared or not.

**Figure 2.**
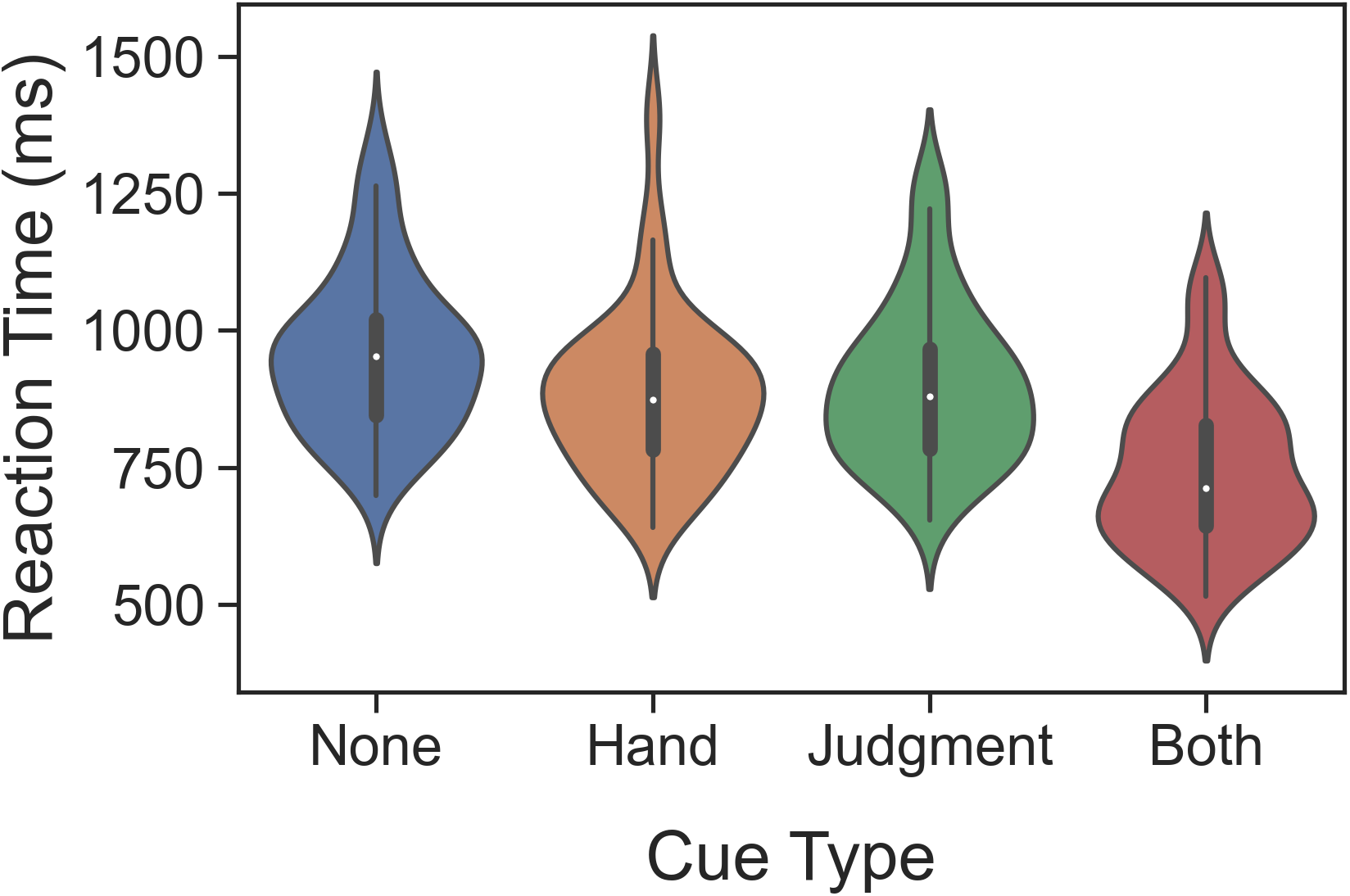
Reaction time (RT) as a function of cue type. Subject-level variability is presented as violin plots within each condition. The center white dot is the mean RT across subjects for that condition. Thick center lines in each violin plot show the 50% inter-quartile range (IQR) and thin lines extend to 1.5*IQR.

**Table 3.** Pairwise Comparisons of Main Effect of Cue Type on RT

| Cue Type Pair |  |  | Mean (ms) | S.E. (ms) | *p* |
| --- | --- | --- | --- | --- | --- |
| Both | — | Response Hand | -141 | ± 6 | < .001* |
|  | — | Judgment Type | -157 | ± 6 | < .001* |
|  | — | None | -215 | ± 10 | < .001* |
| Response Hand | — | Judgment Type | -16 | ± 7 | 0.148 |
|  | — | None | -74 | ± 8 | < .001* |
| Judgment Type | — | Below | -57 | ± 7 | < .001* |

Note: S.E.: standard error.

There were also interactions between cue type and two other factors: 1) judgment domain, *F*(3, 48) = 8.69, *p* < .001, *η_p_^2^* = .35, indicating that RT differences between cue types further depended on whether participants were making a spatial or object-based judgment; and 2) between-subjects congruency, *F*(3, 48) = 5.08, *p* = .004, *η_p_^2^* = .24. Participants benefitted more from the cue in the case of 1) object-based versus spatial judgments and 2) congruent versus incongruent mapping structures. In both cases, these interaction effects did not impact the qualitative pattern of the influence of the amount of cue information on RT.

### Regional Activity Results

#### ROI distribution

As described in detail in the methods, we defined our ROIs for this analysis by generating a grid of spherical ROIs centered around the centroids from a previously-defined atlas (Power et al., 2011) and masking them to our whole-brain task-versus-baseline contrast. Ten ROIs were generated from this analysis, shown overlaid on the task-versus-baseline contrast in Figure 3. Seven of these ROIs were located in left LFC, making these seven ROIs and their relative positioning of particular interest for relating the current results to previous investigations of rostro-caudal organization in the lateral frontal cortex (Badre, 2008; Badre & D’Esposito, 2007; Koechlin & Summerfield, 2007; Nee & D’Esposito, 2016). Four were located in caudal LFC, arranged roughly along the dorsoventral axis and curving rostrally toward the ventral end. The remaining three were positioned rostral to these and were distributed rostro-caudally along the inferior frontal gyrus.

**Figure 3.**
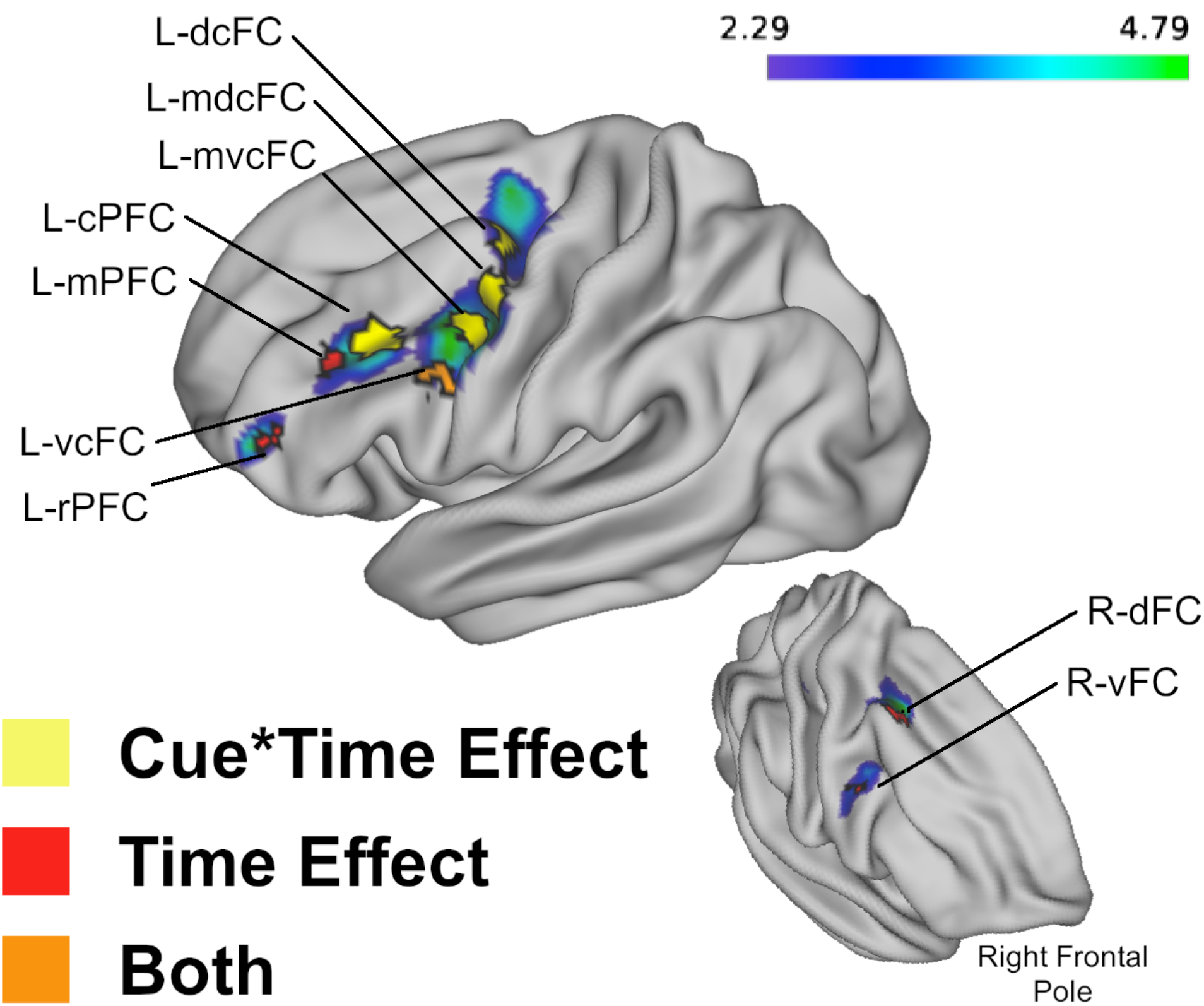
Topographic distribution of frontal ROIs and their associated effects overlaid on task-versus-baseline whole-brain contrast. Left caudal ROIs demonstrated an interaction effect between cue type and time point. Left rostral LFC and right LFC/subcortical ROIs (putamen not shown) demonstrated a main effect of time point. L-vcFC was the only ROI to show both effects. Activity and ROIs have been projected onto surface rendering from volumetric space for ease of visualizing their relative positions in the LFC.

Two ROIs were located in right LFC; one each in caudal dorsal and ventral LFC. Finally, one ROI was located subcortically in the putamen (not illustrated).

#### ROI Analysis

We conducted ten separate four-way ANOVAs, one within each ROI, yielding a total of 150 main and interaction effects across all ROIs. While the factors of judgment domain and response hand were involved in some effects, they were few (9 terms across 5 ROIs) and with small effect sizes (Mean *η_p_^2^*: 0.205; Range: [0.191 : 0.400]). These effects are summarized in Table 4 for reference. To focus on the main research questions of this study, we focus on the main effects of cue type and time point as well as their interaction in the rest of the results of this report.

**Table 4.** Summary of ROI effects involving response hand and judgment domain factors.

| Effect | Regions | *F* (*df*) | *p* | $\eta_p^2$ |
| --- | --- | --- | --- | --- |
| JDxTP | L-mdcFC | 5.055 | 0.038 | 0.229 |
|  | L-cPFC | 11.322 | 0.004 | 0.400 |
|  | L-mPFC | 6.496 | 0.021 | 0.276 |
|  | R--vFC | 11.029 | 0.004 | 0.395 |
| CTxJD | L-mvcFC | 6.192 | <.001 | 0.267 |
|  | L-mPFC | 4.017 | 0.012 | 0.191 |
| JDxRH | L-cPFC | 6.363 | 0.022 | 0.272 |
| CTxRHxTP | L-mPFC | 3.412 | 0.024 | 0.167 |
| CTxJDxRH | Putamen | 4.019 | 0.012 | 0.191 |

Note: CT: cue type factor; JD: judgment domain factor; RH: response hand factor; TP: time point factor; L-: left; R-: right; FC: frontal cortex; PFC: prefrontal cortex; d: dorsal; v: ventral; c: caudal; m: mid; d: dorsal.

Four ROIs demonstrated a significant main effect of cue type, including the left dorsal caudal frontal cortex (L-dcFC; *F*(3,51) = 6.18, *p* = .001, *η_p_^2^* = .27), left mid-dorsal caudal frontal cortex (L-mdcFC; *F*(3,51) = 3.52, *p* = .021, *η_p_^2^* = .17), left mid-ventral caudal frontal cortex (L-mvcFC; *F*(3,51) = 4.48, *p* = .007, *η_p_^2^* = .21), and left caudal prefrontal cortex (L-cPFC; *F*(3,51) = 4.86, *p* = .005, *η_p_^2^* = .22). Activity in these ROIs increased with the amount of information provided by the cue. This effect appeared to be primarily driven by lower activity to noninformative cues overall relative to the informative cues.

All four of these ROIs also demonstrated an interaction between cue type and time point (L-dcFC: *F*(3,51) = 15.34, *p* < .001, *η_p_^2^* = .47; L-mdcFC: *F*(3,51) = 7.67, *p* < .001, *η_p_^2^* = .31; L-mvcFC ( *F*(3,51) = 9.83, *p* < .001, *η_p_^2^* = .37; L-cPFC: *F*(2.341,39.803) = 5.88, *p* = .004, *η_p_^2^* = .26) as well as the left ventral caudal frontal cortex (L-vcFC *F*(3,51) = 9.46, *p* < .001, *η_p_^2^* = .36), indicating that the activity differences in these ROIs by the amount of information contained in the cue further varied as a function of time point within the trial. That is, the cue time point showed generally increasing activity in these ROIs with more cue information, whereas the stimulus time point showed generally decreasing activity with more cue information.

Of these ROIs, only the L-vcFC also exhibited a main effect of time (*F*(1,17) = 9.88, *p* = .006, *η_p_^2^* = .37), indicating that activity in this region was higher at the stimulus overall than at the cue. The remaining ROIs consistently and only demonstrated this main effect of time, including the left middle prefrontal cortex (L-mPFC: *F*(1,17) = 6.52, *p* = .021, *η_p_^2^* = .28); left rostral prefrontal cortex (L-rPFC: *F*(1,17) = 19.97, *p* < .001, *η_p_^2^* = .54); right ventral frontal cortex (R-vFC; *F*(1,17) = 20.86, *p* < .001, *η_p_^2^* = .55); right dorsal frontal cortex (R-dFC: *F*(1,17) = 10.43, *p* = .005, *η_p_^2^* = .38); and putamen (*F*(1,17) = 23.93, *p* < .001, *η_p_^2^* = .58). Activity in these ROIs was significantly higher during the stimulus phase than the cue phase of the trial.

Generally, these results indicate an interaction between cue type and time point or a main effect of time, but not both (with the exception of L-vcFC). Based on these results, we categorized our ROIs by their pattern of effects, with L-dcFC, L-mdcFC, L-mvcFC, and L-cPFC falling in the “Cue*Time” category, and L-mPFC, L-rPFC, R-vFC, R-dFC, and putamen falling in the “Time” category. The left LFC ROIs illustrated in Figure 3 are colored by their effect category; of particular interest, left-hemisphere regions in the “Cue*Time” category were located more caudally, whereas those in the “Time” category were located more rostrally. In other words, regions whose activity differences between time points were dependent on the amount of information contained in the cue were located in areas of frontal cortex associated with lower task abstractness, while those whose activity differences were independent of cue information were located in areas associated with higher task abstractness. Table 5 gives descriptive information about each ROI and summarizes the effects of interest found for each, described in detail below. To illustrate the difference in effects between these two groups, the PSC from baseline as a function of cue type and time point has been plotted in Figure 4 for each ROI from each group. We also present the whole-brain contrasts of each cue level and time point combination versus baseline in Figure 5.

**Table 5.**
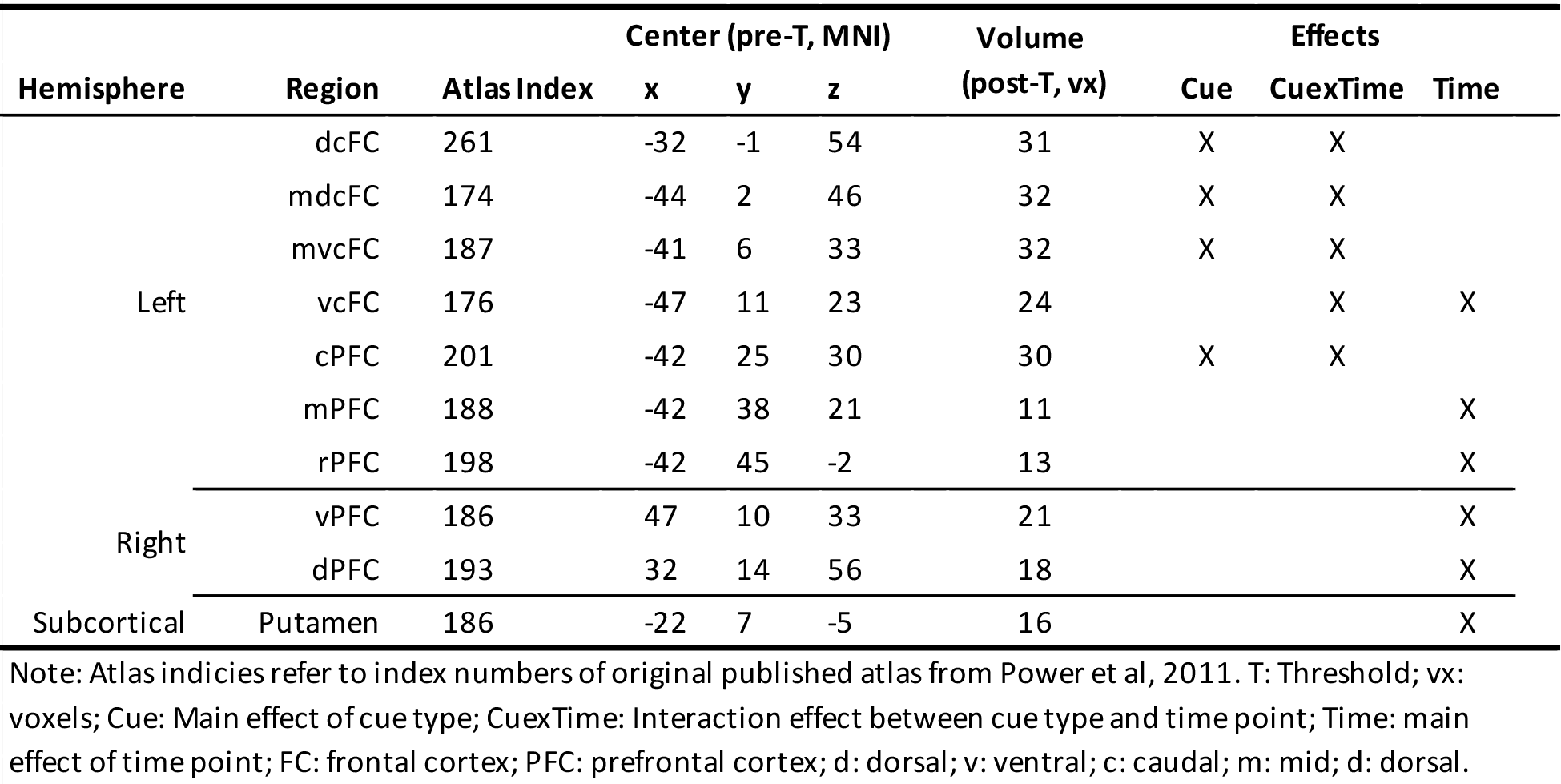
Summary of ROI characteristics and effects.

| Hemisphere | Region | Atlas Index | Center (pre-T, MNI) |  |  | Volume<br>(post-T, vx) | Effects |  |  |
| --- | --- | --- | --- | --- | --- | --- | --- | --- | --- |
|  |  |  | x | y | z |  | Cue | CuexTime | Time |
| Left | dcFC | 261 | -32 | -1 | 54 | 31 | X | X |  |
|  | mdcFC | 174 | -44 | 2 | 46 | 32 | X | X |  |
|  | mvcFC | 187 | -41 | 6 | 33 | 32 | X | X |  |
|  | vcFC | 176 | -47 | 11 | 23 | 24 |  | X | X |
|  | cPFC | 201 | -42 | 25 | 30 | 30 | X | X |  |
|  | mPFC | 188 | -42 | 38 | 21 | 11 |  |  | X |
|  | rPFC | 198 | -42 | 45 | -2 | 13 |  |  | X |
| Right | vPFC | 186 | 47 | 10 | 33 | 21 |  |  | X |
|  | dPFC | 193 | 32 | 14 | 56 | 18 |  |  | X |
| Subcortical | Putamen | 186 | -22 | 7 | -5 | 16 |  |  | X |

Note: Atlas indices refer to index numbers of original published atlas from Power et al, 2011. T: Threshold; vx: voxels; Cue: Main effect of cue type; CuexTime: Interaction effect between cue type and time point; Time: main effect of time point; FC: frontal cortex; PFC: prefrontal cortex; d: dorsal; v: ventral; c: caudal; m: mid; d: dorsal.

**Figure 4.**
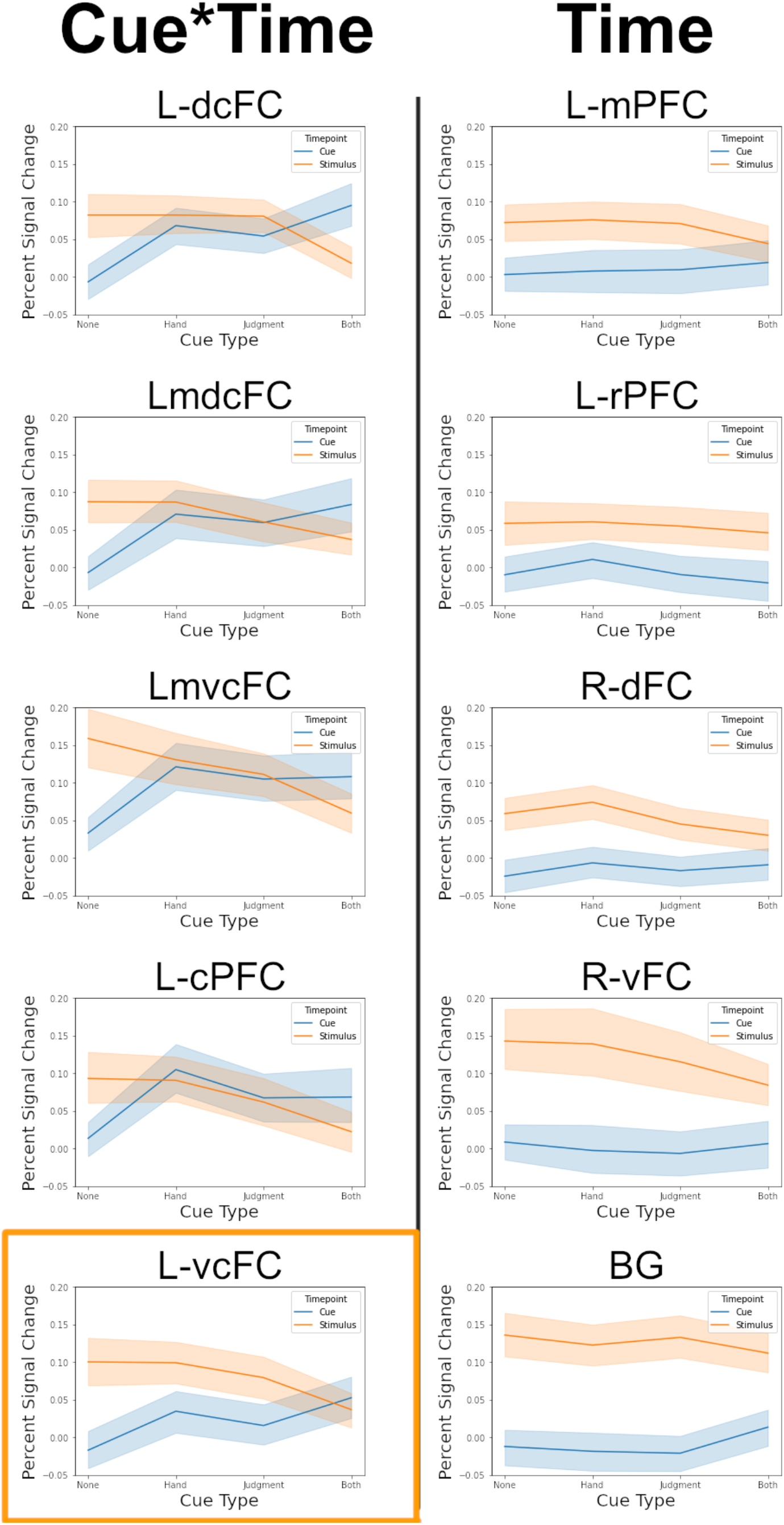
Effects of cue type and time point on PSC in the Cue*Time and Time group ROIs as measured by the BOLD signal. The Cue*Time group demonstrated an interaction between cue type and time point such that activity in these ROIs increased with increasing cue information at the cue time point and decreased with increasing cue information at the stimulus time point. The Time group demonstrated only a main effect of time, such that activity was generally higher at the stimulus than the cue across cue types. L-vcFC, outlined in orange, showed both effects of interest. Subject-level variability is presented as 95% confidence intervals within each condition.

**Figure 5.**
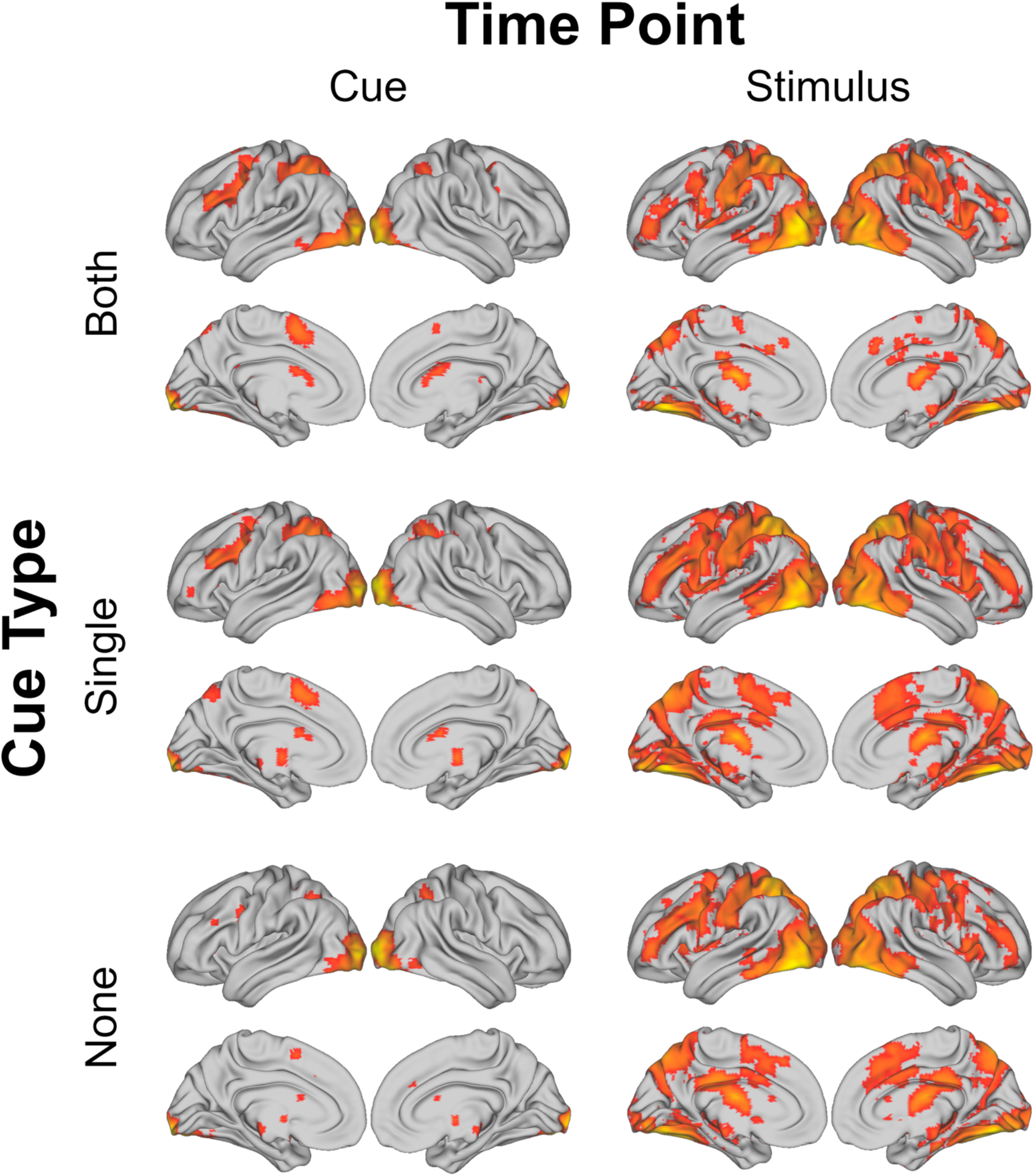
Whole-brain contrasts versus baseline by cue type and time point. All activation maps are shown as PSC thresholded to FDR-corrected *q* = .05. Activity has been projected onto surface rendering from volumetric space for ease of visualizing the distribution across the LFC and other brain areas.

### Exploratory Follow-Up Analysis

Given our observations of the results of our initial analyses, we followed up with an exploratory investigation of the background connectivity among our task-relevant regions and with regions known to be involved in networks supporting cognitive control at different timescales (Dosenbach et al., 2007, 2008). That our two categories of regions were distinguished by two distinct effect patterns involving time was intriguingly reminiscent of the time-based distinctions between the frontoparietal (FP) and cinguloopercular (CO) networks described by the Dual-Networks model of cognitive control. More specifically, the FP network is hypothesized to be involved in moment-to-moment task processing, while the CO network handles tonic task maintenance. These functions were originally supported by evidence that activity in CO regions varied with block level changes in the task rules, while activity in FP regions varied from trial to trial as a function of trial condition. However, these analyses were conducted at the whole-trial level, leaving open the question of how each network’s activity might vary within-trial. One hypothesis in the context of the current experiment is that FP regions support the integration of event information across a trial in pursuit of response selection as it is presented, while CO regions activate on execution of the task regardless of the trial-specific condition to reinforce the currently maintained task representation and evaluate the results. These two functions might then be represented by our two groups of regions in the current study – “Cue*Time” regions supporting the event-related task processing of the FP network, and “Time” regions supporting the task maintenance and evaluation of the CO network. Accordingly, the “Cue*Time” regions were located in areas typically associated with the FP network, while the “Time” regions were in areas associated with the CO network or near the boundary of the two networks.

### Analysis Method

To define ROIs for the dual-networks model, we calculated 6mm spherical regions centered on the point coordinates reported for each region identified in the original model (MNI-translated), generating 18 ROIs. We excluded any ROIs from the dual-networks model that overlapped with any of the ROIs defined in our task data at any point, which removed the left and right frontal cortex regions, the left dorsolateral prefrontal cortex region, and the superior dorsal anterior cingulate/medial superior frontal cortex region; this left left 14 ROIs, for a total of 34 ROIs across both the dual-networks model and the current task. For the primary analyses, we excluded the correlations for the L-vcFC region in this study, as it demonstrated both “Cue*Time”- and “Time”-based effects; the connectivity of this region is presented separately.

Next, we extracted a z-scored time-series correlation matrix across all 34 ROIs using AFNI’s 3dNetCorr function (-fish_z option) applied to the residual time-series generated by the first-level GLM analysis for each subject. This residual time-series which is used as an estimate of “background” activity that has had task-related effects removed. The use of this background activity during correlation gives a measure of latent connectivity between regions that has been isolated from concurrent activation due to mutual involvement in the task (Otten et al., 2002; Tompary et al., 2018). Correlations were calculated across the concatenated time-series of all runs.

Within each resulting correlation matrix, the ROIs were labeled according to their membership in the FP or CO network (in the case of the dual-networks regions) or their characteristic effect pattern (in the case of the task-related ROIs). These labels could then be used to define how pairs of correlated regions contributed to different within- and between-network connectivity values. For example, the correlation value of two regions from the FP network would contribute to the value of the within-FP connectivity, whereas the correlation value of a region from the FP network with a region from the CO network would contribute to the value of the FP to CO between-network connectivity. Correlation values across ROIs from each network pairing were averaged within each subject to yield a network-level correlation matrix for each subject.

Three comparisons were made between values within this network-level correlation matrix. First, we conducted two paired t-tests between the FP-CO between-network connectivity values and each of the FP and CO within-network connectivity values to confirm that the current data demonstrated higher connectivity within versus between networks as has been seen in prior literature. We next replicated these paired t-tests for the two sets of task-related ROIs to explore whether the differences in activity-related effects were likewise associated with correlations reflecting a dual-network structure. Finally, to explore how each set of task-related ROIs were directly connected to the FP and CO networks, we conducted a 2×2 repeated measures ANOVA with factors for network (FP vs CO) and effect pattern (Cue*Time vs Time). If the time-based effect patterns that distinguished our two categories of regions are reflective of the distinct timescales at which the FP and CO networks instantiate cognitive control, the connectivities within and between regions of our two effect categories should mirror the connectivity patterns seen in regions of the FP and CO networks, and the connectivity between regions of our two effect categories and those of the dual-networks model should show an interaction between effect category and connecting network. Finally, we conducted paired *t*-tests on the correlation values between L-vcFC and each of the sets of regions for the FP and CO networks as well as the “Cue*Time” and “Time” groups.

### Follow-Up Results

The FP and CO regions from the dual-networks model (Dosenbach et al., 2007) both demonstrated significantly higher within-network connectivity relative to their between-network connectivity across the session (FP: *t*(17) = 9.388, *p* < .001, Cohen’s *d* = 2.213; CO: *t*(17) = 9.857, *p* < .001, Cohen’s *d* = 2.323). Connectivity also varied in the two sets of regions identified above. However, while the “Cue*Time” regions showed greater connectivity to each other than with the “Time” regions (*t*(17) = 4.862, *p* < .001, Cohen’s *d* = 1.146) as expected, the “Time” regions showed the opposite pattern (*t*(17) = −3.011, *p* = .008, Cohen’s *d* = −.710). That is, the “Time” regions were more strongly connected to the other regions identified in the task than they were to each other. These results are contrasted in Figure 6.A.

**Figure 6.**
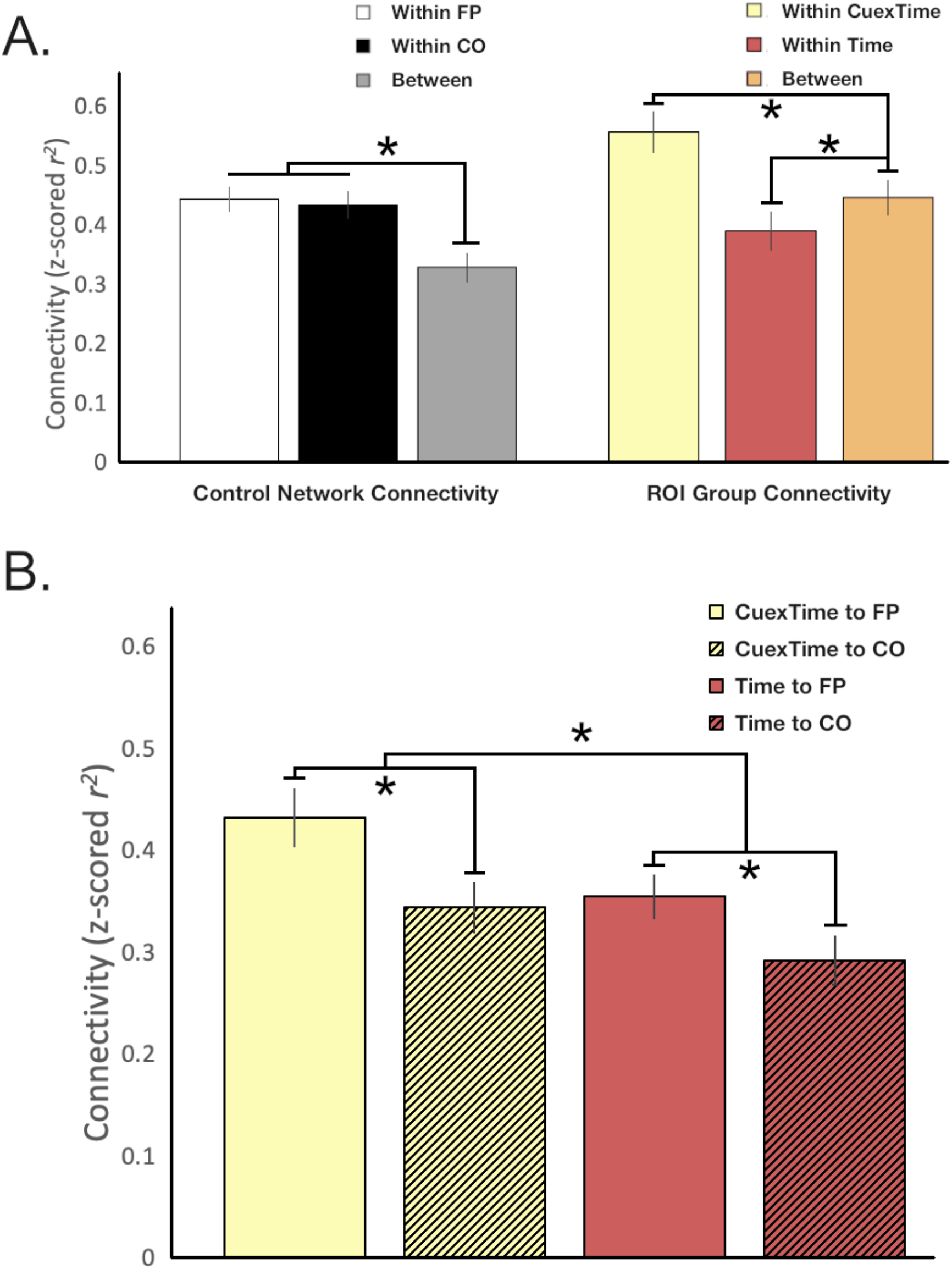
Results of exploratory connectivity analyses. A) Comparison of connectivity patterns seen in known cognitive control networks with those seen in the Cue*Time and Time groups of ROIs from the current study. Between = between-network/group connectivity. B) Connectivity between the ROIs from the current study and each of the two cognitive control networks. Error bars in all graphs represent the standard error within conditions.

The ANOVA assessing the “Cue*Time” and “Time” regions’ connectivities with FP and CO regions further revealed main effects of network (*F*(1,17) = 46.600, *p* < .001, *η_p_^2^* = .733) and of effect category (*F*(1,17) = 28.659, *p* < .001, *η_p_^2^* = .628). Overall, our regions demonstrated greater connectivity with FP regions than CO regions, and “Cue*Time” regions had stronger connectivity than “Time regions” across both networks. The interaction between these factors was not significant (*F*(1,17) = 2.421, *p* = .125). These results are illustrated in Figure 6.B. Taken together, these results indicate that both our “Cue*Time” and “Time” regions are more strongly associated with the FP network than the CO network, and that our “Cue*Time” regions are especially tightly coupled with regions associated with moment-to-moment processing.

The connectivity between the L-vcFC region and each set of regions (FP, CO, “Cue*Time”, and “Time”) is presented in Table Y. Paired *t*-tests demonstrated that the L-vcFC was more strongly connected to the “Cue*Time” regions than “Time” regions (ΔPSC = 9.3 ± 2.1%, *t*(17) = 4.435, *p* < .001), and was equally connected to both the FP and CO regions (ΔPSC = 1.0 ± 2.3%, *t*(17) = 0.406, *p* = .690). Additionally, the connectivity between the L-vcFC and both the “Cue*Time” and “Time” regions were generally higher than that between the L-vcFC and either the FP or CO regions (ΔPSCs between 10.0 ± 1.8% and 20.3 ± 2.6%, *t*(17) between 5.664 and 11.363, all *p* < .001).

## Discussion

The current study is, to our knowledge, the first investigation of how nested task set representations may be implemented by the cognitive control system using a pre-cueing procedure to separate task processing across events in time while conferring different amounts of cue information from trial to trial. Through this design, we aimed to determine the neural correlates of task processing and task representation and the potential overlap between them. Regions that support preparation and execution processes should show activity that varies from one time point in the trial to another; those that are involved in task representation should show activity that varies with the amount of information contained in the cue. First, we asked whether task processing and task representation are distinct aspects of cognitive control or are part of the same underlying control mechanism. In the former case, regions would be sensitive to only one factor or the other (i.e., time point and cue type), whereas in the latter some or all regions should demonstrate an interaction between them. This also gave us the opportunity to verify that cues produce performance benefits even when they do not include any information about the upcoming response. Next, we asked whether cue-sensitive regions showed changes in cue-related activity that reflected preparation of sets of different sizes or reduction of different amounts of uncertainty, based on whether they showed decreasing or increasing activity at the cue time point. Finally, based on differences in the effect patterns that we observed in left caudal frontal regions relative to other task related regions, we further conducted an exploratory analysis that aimed to relate these different sets of effects to network-level connectivity.

Our data indicate that amount of cue information and time point did indeed interact in a number of our task-relevant regions (viz., L-dcFC, L-mdcFC, L-mvcFC, L-vcFC, and L-cPFC; see Figure 3), clearly indicating a shared mechanism of task processing and representation. The more of the task that is resolved at the cue, the less needs to be done when the stimulus appears. Interestingly, all regions that demonstrated an effect of cue information also showed this interaction, while most regions demonstrating a main effect of time point did not (viz., L-mPFC, L-rPFC, R-dFC, R-vFC, putamen, with the exception of L-vcFC), suggesting they may be separately involved in task execution. Thus, preparation mechanisms appear to be inextricably interlinked with task representation, uniquely supported by left caudal frontal regions; the L-vcFC, showing effects associated with both groups of regions, may uniquely be involved in bridging between these preparatory processes and downstream execution processes.

Of our task-related regions, those demonstrating an interaction between cue information and time point were restricted to left-lateralized mid-caudal to caudal LFC. The pattern of activity seen for different types of cues was relatively similar across these regions. At the cue, regions generally showed the least activity for a noninformative cue, roughly equal levels of activity for each of the cues giving one piece of information, and the most activity for cues for both pieces of information; this pattern was reversed at the stimulus. As noted above, this mirrors the stair-step effect seen in the behavioral data, suggesting that this activity may be a neural correlate of the cue benefit itself. Activity in these regions may reflect the integration of new cue information with the existing task representation to reduce uncertainty about the upcoming task at the cue, where the processing of more information demands greater activity. At the stimulus, then, processing demand in these regions is likewise greater when more of the task remains to be specified. Lower cue information thus causes increased demand at the stimulus that may require longer processing time, resulting in the stair-step effect seen both in the current reaction time data and likewise in previous behavioral studies (Rosenbaum, 1980). Previous research on reward motivation at different levels of task representation used a psychophysiological interaction (PPI) analysis to demonstrate a positive relationship between frontal activity and reward-based performance benefits that was specific to task representation level (Bahlmann et al., 2015). This may hint at a similar relationship for frontal activity and cue benefits. While the current study does not have the power to assess the relationships between our frontal activation and cue benefit at each task representation level, future work using the hierarchical pre-cue design could remove noninformative cues, to allow for a direct, more powerful comparison between two different cue levels.

Beyond the relationship between cue benefits and cue information, the pattern of activity seen in these regions also indicates that the level of activity at each task event in the pre-cueing procedure is related to the amount of information being simultaneously integrated, or alternatively the amount of uncertainty being resolved at that time point, rather than the size of the task set being prepared at the time of the event. At the cue, participants showed lower activity with less cue information. This is perhaps expected for noninformative cues, which have previously shown lower activity in frontal cortex relative to informative cues in standard pre-cueing procedures (e.g., Cookson et al., 2016). More interesting is that this pattern is also seen between different levels of informative cues. In our design, cues for one piece of task information reduced the task set to half the total possible decision options (i.e., two 2-alternative forced choice tasks), while cues for both pieces reduced the task to a single 2-alternative forced choice task. That there is greater frontal activity at the cue for cues giving reduce uncertainty of the number of responses rather than to activate smaller sets. This is further supported by the reversal of the pattern at the stimulus; as participants have processed more of the task a priori, there is less remaining at the stimulus, so cues for both pieces of information have the least activity.

A third intriguing observation from these activity patterns is that they do not appear to vary with rostro-caudal position. The regions in left LFC demonstrating an interaction between cue information and time point roughly align with regions previously associated with the lower two levels of a policy abstraction task (Badre & D’Esposito, 2007), suggesting that a two-level task set representation was employed by participants in this study. The question then remains what aspects of the task structure are represented at each level, and how the cue information influences processing at each level. That the activity in the caudal and mid-caudal regions in this study followed the same patterns of activity across cue types indicates that cues were informative at both levels of representation. This may be a reflection of the nature and flexibility of task abstraction in the brain. Coding of tasks in LFC is highly flexible (Cole et al., 2014; Stokes et al., 2017), and tasks can be represented with a wide variety of task features (Badre & D’Esposito, 2007; Cookson et al., 2019; Huang & Awh, 2018). This includes cue information, even if it is presented during distinct time points (Grant et al., 2020; Hazeltine et al., 2011). Because the cue is always presented first in this experiment, it is possible that participants represented the highest level of the task as a distinction of the amount of information contained in the cue. Then, based on the type of cue presented, they could then determine what kind of reduction, if any, they would be able to make in the upcoming task, and prepare accordingly. Future research should explore this type of content-independent meta-task representation and its relationship to policy abstraction and the rostro-caudal axis.

Our behavioral results may provide some insight into how these mechanisms interact. The current results demonstrated a stair-step effect as a function of the amount of information provided by the cue, such that participants responded more quickly and with higher accuracy based on how much information they were given, independent of what those pieces of information were. This mirrors similar effects seen for specification of individual components of a motor response (Rosenbaum, 1980), despite the use of cues in the current experiment that expressly could not be used to specify a motor component of the task. Here, single-level cues for both judgment domain and response hand showed not only similar patterns of behavioral benefits. But similar patterns of activity in Cue*Time regions as well. Both judgment domain and response hand cues allowed participants to reduce the size of the relevant task by half, but only the response hand cue indicated a dimension of the response to be prepared. That the patterns for both cue types were the same suggests that participants were indeed using the cue information to prepare for the task at the task set level and not simply activating responses.

However, preparation of motor responses and more abstract task sets may involve the same general mechanism implemented at different levels of representation. Badre and Frank (2012; see also Frank & Badre, 2012, for computational model) have previously demonstrated a similar distribution of a reinforcement learning mechanism across distinct task levels. This mechanism is supported by progressive cortico-subcortical loops between frontal regions distributed along the rostro-caudal axis and similarly organized subregions of the basal ganglia. Each loop implements gating mechanisms at a given level of representation to learn probabilistic associations at that task level; the results of that selection process then cascade from the top levels of abstraction to the most concrete to coordinate across levels. Gating mechanisms are critical for cognitive control (Chiew & Braver, 2017; Ott & Nieder, 2019) and specifically for task preparation (Ruge & Braver, 2007); this distributed gating mechanism, then, likely also supports selection between task representations at each level as a function of the representational content maintained in LFC within each level of cortico-subcortical loops along the rostro-caudal axis. In the current task, then, participants may implement cognitive control through a two-level gating mechanism, selecting between abstract task sets within the task representation as a whole at one level and preparing specific stimulus-response sets at the other.

Throughout this report, we have discussed participants’ use of nested, abstracted task sets in their representation and execution of our task. These are closely related to ideas of representational and processing hierarchies that have been proposed to underlie cognitive control (Badre & D’Esposito, 2007; Koechlin & Summerfield, 2007; Nakayama et al., 2016; Nee & D’Esposito, 2016). Indeed, the task file hypothesis on which the current study bases its predictions itself proposes a hierarchical structure for task sets across different levels of representation. Typical studies investigating hierarchical processing and its neural correlates use tasks that use context-driven policy abstraction – that is, tasks in which higher-order rules dictate the relevant lower-level rules and mappings and whose information must be integrated with the rest of the task to select a response (see Badre, 2008). The current task does not use this type of hierarchical mapping – participants can complete the task regardless of the information contained in the cue with only the information presented with the two target stimuli. Nonetheless, they do appear to represent the task with a nested task set structure that can divide the task abstractly along judgment domain (spatial/object) and/or response hand (left/right) dimensions. It is entirely possible that this nested task set structure is implemented via a hierarchical mechanism, although it would not be necessary in this task. In contrast to early theories of the rostro-caudal hierarchy that hypothesized specific brain areas associated with progressively abstracted representations, Badre and Nee (2018) have proposed a more flexible hierarchy in which the LFC is grossly subdivided into three general zones supporting “sensory-motor”, “contextual”, and “schematic” control from caudal to rostral, respectively. In particular, contextual control areas support a wide range of high-level action representation and processing, ranging from simple rule associations to full task representation. Hommel (2021) has proposed GOALIATH, a theory of goal-directed action selection that would allow these contextual control areas to flexibly represent and implement task-relevant actions without prescriptivist definitions of the role of individual regions within this zone. Our results support such an interpretation; our “Cue*Time” regions indeed fall within the “contextual control” zone described by Badre and Nee, and regions within this area show similar patterns of activity to the cue regardless of position, as they would if they were being flexibly recruited to represent and prepare the task from trial to trial.

As a final point, there is a difference between the groups of regions that were involved in representational aspects of task processing, evidenced by an interaction between cue information and time point (”Cue*Time” group) and those that were involved in other aspects of task processing, exhibited by a main effect of time point (”Time” group). Specifically, the rostro-caudal distribution of these effect patterns in left LFC. Our “Cue*Time” regions are located in more caudal LFC whereas “Time” regions are located more rostrally. This is similar to the distribution of distinct sets of regions involved in two different networks related to cognitive control, as described by the Dual Networks model (Dosenbach et al., 2007, 2008). Our “Cue*Time” regions are located in areas similar to those regions in the FP network of the dual-networks model, while our “Time” regions are in areas similar to those regions of the CO network. Moreover, the FP network is hypothesized to facilitate moment-to-moment processing, whereas the CO network supports tonic maintenance of active representations. This parallels the distinctions in processing and representation effects found for the regions in this study; the interaction between cue information and time-point may capture moment-to-moment processing of cue information, while the main-effect of time may indicate the activation of the task representation during response execution regardless of the specific information being processed. Thus, we conducted an exploratory analysis to assess whether the connectivity of our two groups of regions mirrored that of regions in the FP and CO networks.

However, our background connectivity analysis did not support this hypothesis. Connectivity patterns between FP and CO regions in our data reflected typical network patterns, where connectivity within both networks was greater than connectivity between the two networks (Cohen et al., 2014). In contrast, the connectivity between “Cue*Time” and “Time” regions was in fact higher than the connectivity within our “Time” regions, suggesting that the regions supporting non-representational task processing do not form a cohesive and separate network from those supporting representational task processing. Additionally, both sets of regions were more strongly connected to FP regions than CO regions, with “Cue*Time” regions showing greater connectivity with both networks than “Time” regions. Overall, this generally indicates that the two effect patterns found in this study were not drawn along network boundaries. Rather, it suggests that regions involved in all aspects of task processing in this study are more strongly associated with the FP network than the CO network, i.e., with moment-to-moment processing. In other words, the results presented here are preliminary evidence of a rostro-caudal distinction of regional functions in LFC that appears to occur within the FP network; future research should further explore how these rostro-caudal distinctions fit within the more general functional framework of the dual-networks model.

Taken together, the results of this study demonstrate that task representation is not a separate static function but is one of a subset of active task processing functions subserved by the brain, and that the brain processes tasks from moment to moment by resolving uncertainty for the upcoming action at the level of abstract task sets in addition to direct motor preparation. Furthermore, they begin to draw new parallels between the task preparation and execution literature and the representational abstraction literature, giving us a more complete and cohesive understanding of the implementation of cognitive control in the brain. We hope that making this procedure open-sourced will encourage others to continue to explore this design and learn more about task processing and representation in the brain.

## Acknowledgments

The authors would like to acknowledge Zoey Morton, who managed and conducted the majority of data collection for the present study. We would also like to thank the Georgia Institute of Technology, where the present study was conducted as a part of S.L.C.’s doctoral program. Work for this publication done after January 2019 was made possible by an NIMH postdoctoral fellowship to S.L.C. (5 F32 MH119761-02).

